# Discovery of SARS-CoV-2 antiviral synergy between remdesivir and approved drugs in human lung cells

**DOI:** 10.1101/2020.09.18.302398

**Authors:** Xammy Nguyenla, Eddie Wehri, Erik Van Dis, Scott B. Biering, Livia H. Yamashiro, Julien Stroumza, Claire Dugast-Darzacq, Thomas Graham, Sarah Stanley, Julia Schaletzky

## Abstract

The SARS coronavirus 2 (SARS-CoV-2) has caused an ongoing global pandemic with currently 29 million confirmed cases and close to a million deaths. At this time, there are no FDA-approved vaccines or therapeutics for COVID-19, but Emergency Use Authorization has been granted for remdesivir, a broad-spectrum antiviral nucleoside analog. However, remdesivir is only moderately efficacious against SARS-CoV-2 in the clinic, and improved treatment strategies are urgently needed. To accomplish this goal, we devised a strategy to identify compounds that act synergistically with remdesivir in preventing SARS-CoV-2 replication. We conducted combinatorial high-throughput screening in the presence of submaximal remdesivir concentrations, using a human lung epithelial cell line infected with a clinical isolate of SARS-CoV-2. We identified 20 approved drugs that act synergistically with remdesivir, many with favorable pharmacokinetic and safety profiles. Strongest effects were observed with established antivirals, Hepatitis C virus nonstructural protein 5 A (HCV NS5A) inhibitors velpatasvir and elbasvir. Combination with their partner drugs sofosbuvir and grazoprevir further increased efficacy, increasing remdesivir’s apparent potency 25-fold. We therefore suggest that the FDA-approved Hepatitis C therapeutics Epclusa (velpatasvir/sofosbuvir) and Zepatier (elbasvir/grazoprevir) should be fast-tracked for clinical evaluation in combination with remdesivir to improve treatment of acute SARS-CoV-2 infections.

---

SARS-CoV-2, a positive-sense RNA betacoronavirus, is the causative pathogen for the novel coronavirus disease 2019 (Covid-19) ^1^. Highly transmissible and without a cure or a vaccine, it caused a rapidly spreading global pandemic. SARS-CoV-2 infects human epithelial lung cells via interaction with the ACE2 receptor, followed by virus replication and spread ^2^. Pneumonia and acute respiratory distress can be severe, with alveolar damage and blood clotting abnormalities and unusual large-vessel strokes, often weeks after infection ^3,4^. Sequelae include impaired lung function due to pulmonary fibrosis ^5^, myocardial and neurological events, and the need for a lung transplant ^6-8^. The case fatality rate is currently ∼3.5% in the US, and 4.5% worldwide, indicating that despite improvements in therapy and testing, we cannot effectively treat the disease ^9^.

Best in class and the sole drug with Emergency Use Authorization by the FDA for treatment of Covid-19 is remdesivir (GS-5734), a broad-spectrum antiviral originally discovered to treat Hepatitis C virus (HCV) and Ebola ^10-12^. Remdesivir is a 1′-cyano-substituted adenine C-nucleoside ribose analogue (Nuc), a prodrug that requires intracellular conversion to an active triphosphate metabolite (NTP), which interferes with the activity of viral RNA-dependent RNA-polymerases (RdRp) ^13^. In animal models of Covid-19, robust effects are seen in non-human primates ^14,15^. However, in humans, the median recovery time in a phase III clinical trial treatment group was only reduced from 15 to 11 days ^16^, while in other studies no significant improvement over standard of care was apparent ^17,18^. As a prodrug given intravenously, remdesivir’s pharmacokinetic profile is highly complex with several active metabolites ^15^. As remdesivir is not highly potent, diffusion-driven distribution to the target tissue seems to be limiting efficacy, prompting the evaluation of an inhaled formulation ^19,20^. Alternative approaches to improve remdesivir efficacy are urgently needed.

In antiviral therapy, combination therapies are highly efficacious, safe, and less prone to resistance development ^21,22^. Indeed, the combination therapies Epclusa (velpatasvir/sofosbuvir) and Zapatier (elbasvir/grazoprevir) have transformed Hepatitis C care ^23^. Similar combination approaches for Covid-19 would be highly desirable, as they could increase potency of remdesivir and allow a vastly larger number of patients to be treated with the existing limited stockpile. Thus, we conducted a high-throughput combinatorial screen to identify FDA-approved compounds that act synergistically with remdesivir in blocking SARS-CoV-2 induced cytopathic effect. Among 20 identified compounds that show robust synergy with remdesivir, the largest effects are observed with HCV antivirals velpatasvir and elbasvir, targeting the replication co-factor NS5A. A further increase in synergy was observed upon combining remdesivir with the commercially available HCV combination therapies Epclusa (velpatasvir/sofosbuvir) and Zepatier (elbasvir/grazoprevir). The resulting ∼25-fold increase in remdesivir potency could significantly increase efficacy in the lung target tissue, allow treatment of more than 5 million Covid-19 patients with the doses manufactured in September (currently treating about 230000 patients), and provide an opportunity for developing a more efficacious treatment for Covid-19.

## Combinatorial high-throughput screen for compounds synergistic with remdesivir

We developed a robust high-throughput assay in SARS-CoV-2 infected monkey kidney epithelial Vero E6 and human lung epithelial Calu-3 cells (Figure 1a) ^24^, using a clinical isolate of SARS-CoV-2 virus (SARS-CoV-2 USA-WA1/2020) ^25^. Cells were treated with 40μM compound, infected with SARS-CoV-2, and incubated for 72-96h. We then measured virus-induced cytopathic effect (CPE) by quantifying ATP in viable cells via luminescence. We determined the average EC50 for remdesivir to be 3 +/-0.6μM in Vero-E6 and 0.7+/-0.1μM in Calu 3 cells (Figure ED1), consistent with literature values (Vero-E6 EC50 0.6-11μM, Calu 3 EC50 0.3-1.3μM ^26-28^).

**Figure 1.**
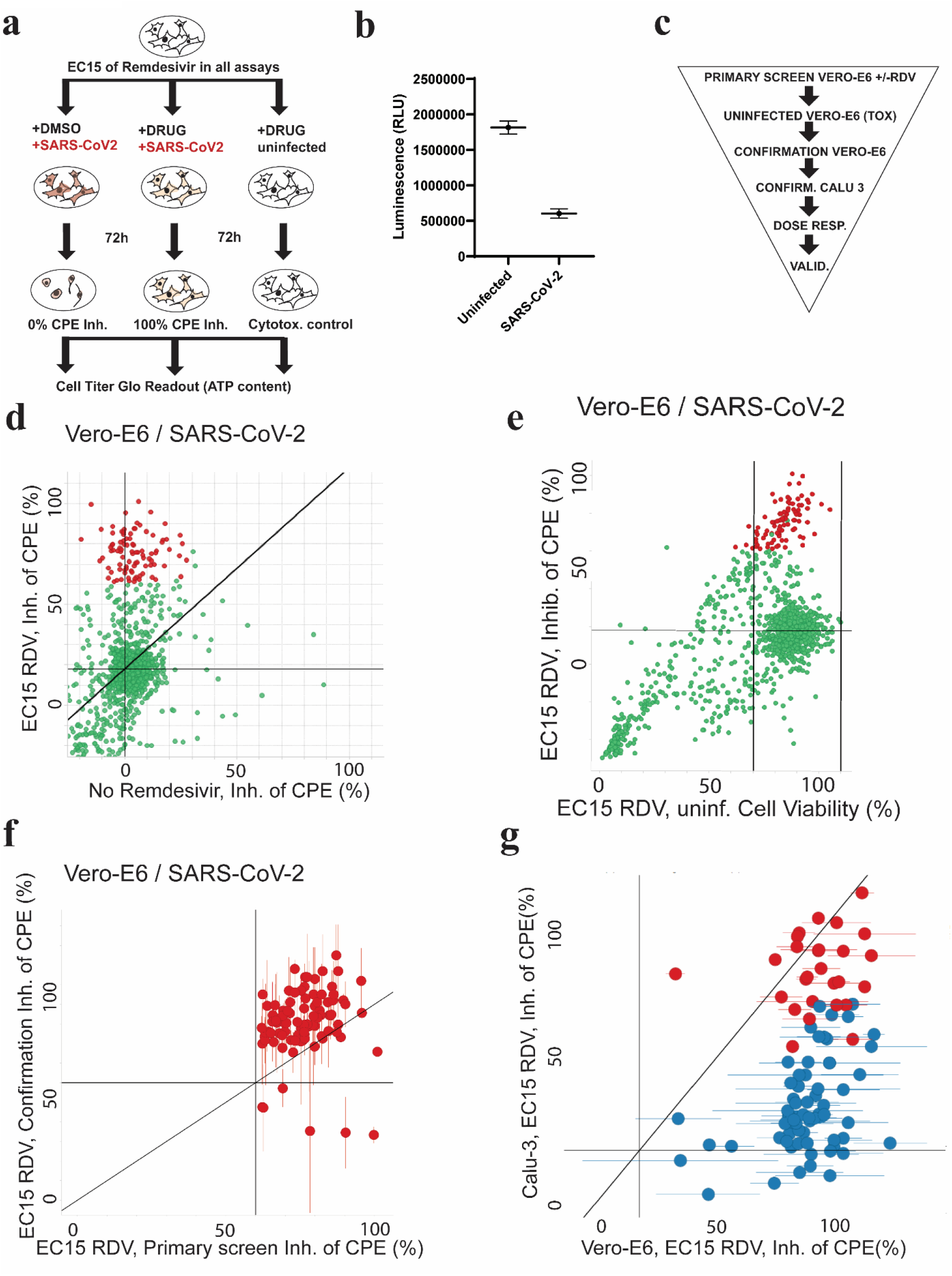
Primary screening results identifying compounds increasing antiviral effects of remdesivir (RDV). **(a)** Assay outline: Vero-E6 cells are added to 384 well plates, treated with DMSO (left panel) or drug (middle panel), infected with SARS-CoV-2 and incubated for 72h to observe cytopathic effect (CPE; left panel). Effective drug treatment inhibits occurrence of CPE (middle panel). CPE is measured by quantifying ATP content in viable cells using a luminescent assay (Cell Titer Glo). The right panel shows the cytotoxicity control, treating cells with drugs but without virus. **(b)** Screening assay performance. Average Luminescence is shown for the Vero E6 primary screen in presence of EC15 of remdesivir (n=144, 24 wells each from 6 screening plates), error bars indicate standard deviation. “Uninfected”: positive control (equivalent to 100% inhibition of CPE), “SARS-CoV-2”: negative control, infected and treated with DMSO (equivalent to 0% inhibition of CPE). Z’=0.63+/-0.04. **(c)** Screening paradigm outline in presence of EC15 of remdesivir. Cells infected with SARS-CoV-2 unless indicated. Dose resp. – Dose response; Valid. – Validation in orthogonal assays. **(d)** Primary screen results for 1200 approved drugs tested in Vero E6 cells infected with SARS-CoV-2 in the absence (x-axis) and presence (y-axis) of EC15 of remdesivir. Inhibition of CPE (%) is shown. The horizontal line indicates the background activity of EC15 of remdesivir (not subtracted). Diagonal line: 1:1 correlation. Red: high priority hits with a cutoff of >60% inhibition of CPE in presence of remdesivir. **(e)** As in (d), but with cell viability data from cytotoxicity control (uninfected) on the x-axis. Vertical line: Cell viability of 70%. **(f)** Confirmation of >95% of high priority hits from (e) after compound cherrypicking; assay conditions as in (e), Vero-E6 cells infected with SARS-CoV-2 in presence of EC15 of remdesivir; x-axis indicates primary screening results (Inhibition of CPE, %), y-axis confirmation results (Inhibition of CPE, %). Horizontal and vertical lines indicate hit progression cutoff from primary screen, diagonal line 1:1 correlation. Error bars indicate standard deviation. **(g)** 26 compounds (red; labeled) are active in both Vero E6 and human lung epithelial Calu-3 cells infected with SARS-CoV-2 in presence of EC15 of remdesivir. Inhibition of CPE (%) is shown on the x-axis for Vero E6, on the y axis for Calu-3 cells.

We then exposed cells to combinations of remdesivir (at concentrations causing 15% inhibition of CPE, EC15) with a library of ∼1200 FDA-approved drugs. A parallel screen in the absence of viral infection assessed compound toxicity (Figure 1a). Importantly, the EC15 concentration of remdesivir (0.3-1μM) is comparable to the serum concentration of the main remdesivir metabolite in plasma (∼0.4 μM) ^14^, indicating that the results of a screen performed under these conditions could be clinically meaningful.

Using these conditions, we were able to precisely measure CPE caused by SARS-CoV-2 infection and achieved an average Z’ of 0.63 +/-0.04 during the primary screen (Figure 1b). The primary screen of 1200 approved drugs identified 90 compounds with antiviral activity exclusively in presence of EC15 remdesivir (Figure 1d, red). None of the hit compounds showed significant toxicity in cells (Figure 1e, red). More than 95% of hit compounds confirmed in an independent Vero E6 assay (Figure 1f). 28 compounds showed strong activity across different cell lines (Vero E6 and Calu-3) in a background of EC15 of remdesivir (Figure 1g, red).

As all tested compounds are approved drugs annotated with their molecular targets, we conducted a gene set enrichment analysis to identify pathways preferentially targeted by hit compounds in the combinatorial screen (Figure 2). We observed a statistically significant enrichment of compounds affecting the corticosteroid pathway (Figure 2, GSEA p=0.0001), as well as for Calcium channel, proton pump and HIV protease modulation (Figure 2, GSEA p= 0.004, 0.003), all of which have been implicated in antiviral effects in the literature ^28-31^. Without remdesivir, none of the mentioned targets were enriched (p>0.1, FDR q-value >0.36) ^32^. This indicates that remdesivir makes SARS-CoV-2 uniquely vulnerable to inhibition of otherwise nonessential targets.

**Figure 2.**
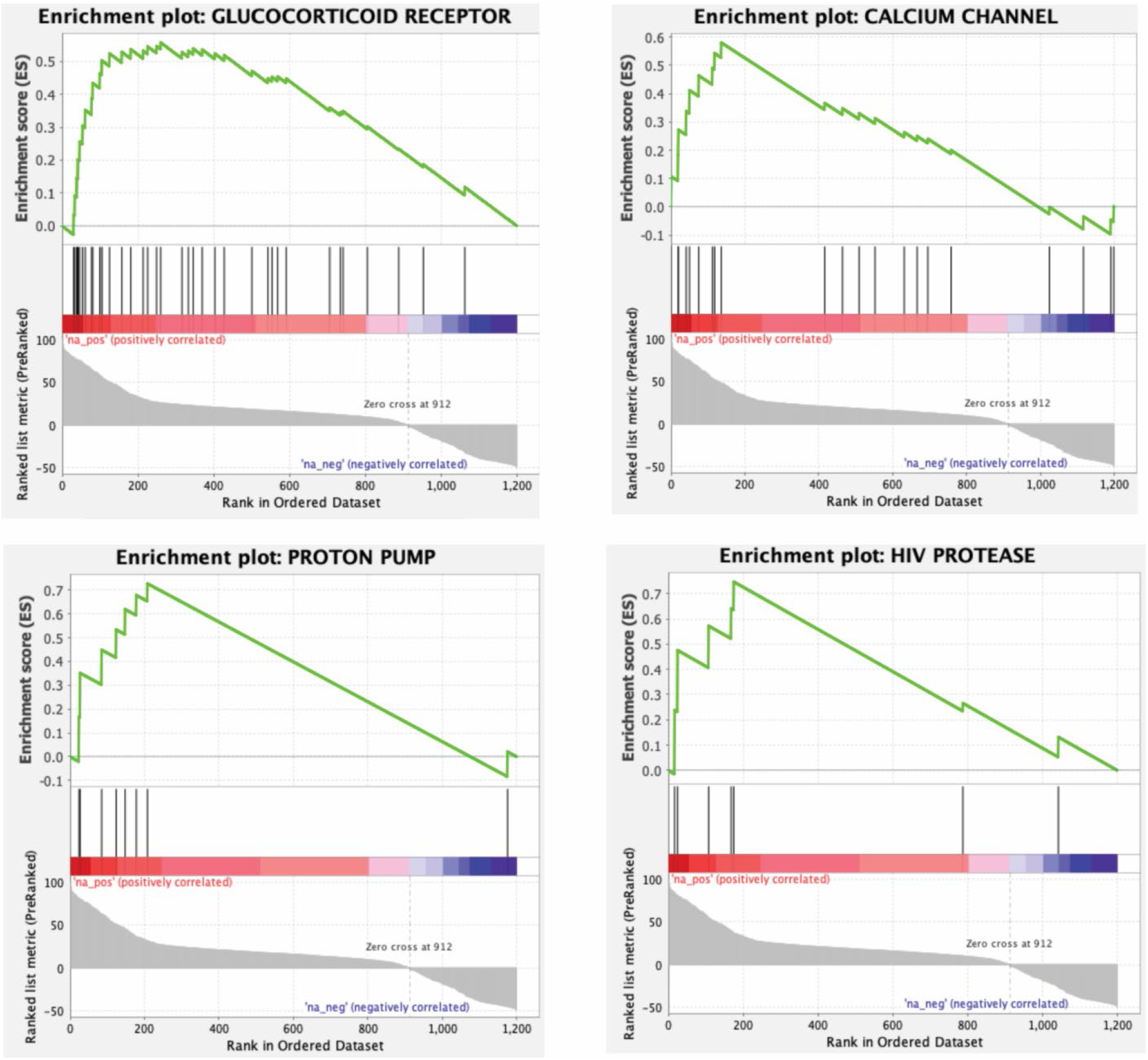
Gene set enrichment analysis of drug targets in combinatorial screen. GSEA enrichment plots provide the distribution of the enrichment score (green line) across compounds annotated to molecular targets (vertical black line), ranked in order of antiviral activity (left to right). The enrichment score (ES) reflects the degree to which a gene set is overrepresented at the top of a ranked list of compounds interacting with the given target. GSEA calculates the ES by walking down the ranked list of compounds interacting with the given target, increasing a running-sum statistic when a gene is in the gene set and decreasing it when it is not. Glucocorticoid receptor (p=0.0001; FDR q-value=0.013), Calcium Channel (p=0.004; FDR q-value=0.086), Proton pump (p=0.003; FDR q-value=0.085) and HIV protease (p=0.007; FDR q-value=0.095) are identified as targets enriched in the hitlist for the synergy screen in background of EC15 of remdesivir.

## Quantitation of synergistic effects with remdesivir in a dose response matrix

To identify the most promising drug combinations for use in the clinic, we conducted a dose-response interaction matrix analysis to quantitatively evaluate the synergy between screen hits and remdesivir. The matrix combined ten concentrations of remdesivir (up to 10μM) with eleven concentrations of each screen hit (up to 40μM), allowing us to test CPE in SARS-CoV-2-infected Calu-3 cells for 110 concentration combinations per remdesivir/compound pair. We then used computational zero interaction potency (ZIP) modeling to quantitatively determine if synergy was present ^33^. The model combines both Loewe additivity and Bliss independence models, systematically assessing drug interaction patterns that may arise in drug combination matrix. In this model, a value of <0 signifies antagonism, 0-10 additive effects, and values >10 show synergy between compound pairs. Strikingly, 20 compounds showed pronounced synergy with remdesivir in counteracting SARS-CoV-2 infection, with maximal ZIP-scores of 29-87 (Figure 3, Figure ED2): velpatasvir, elbasvir, dabrafenib, cilostazol, nimodipine, conivaptan hydrochloride, clobetasol, budesonide, drosiprenone, ezetimibe, ivosidenib, selexipag, meprednisone, nifedipine, omeprazole sulfide, quinapril, rifaximin, telmisartan, valdecoxib and zafirlukast.

**Figure 3.**
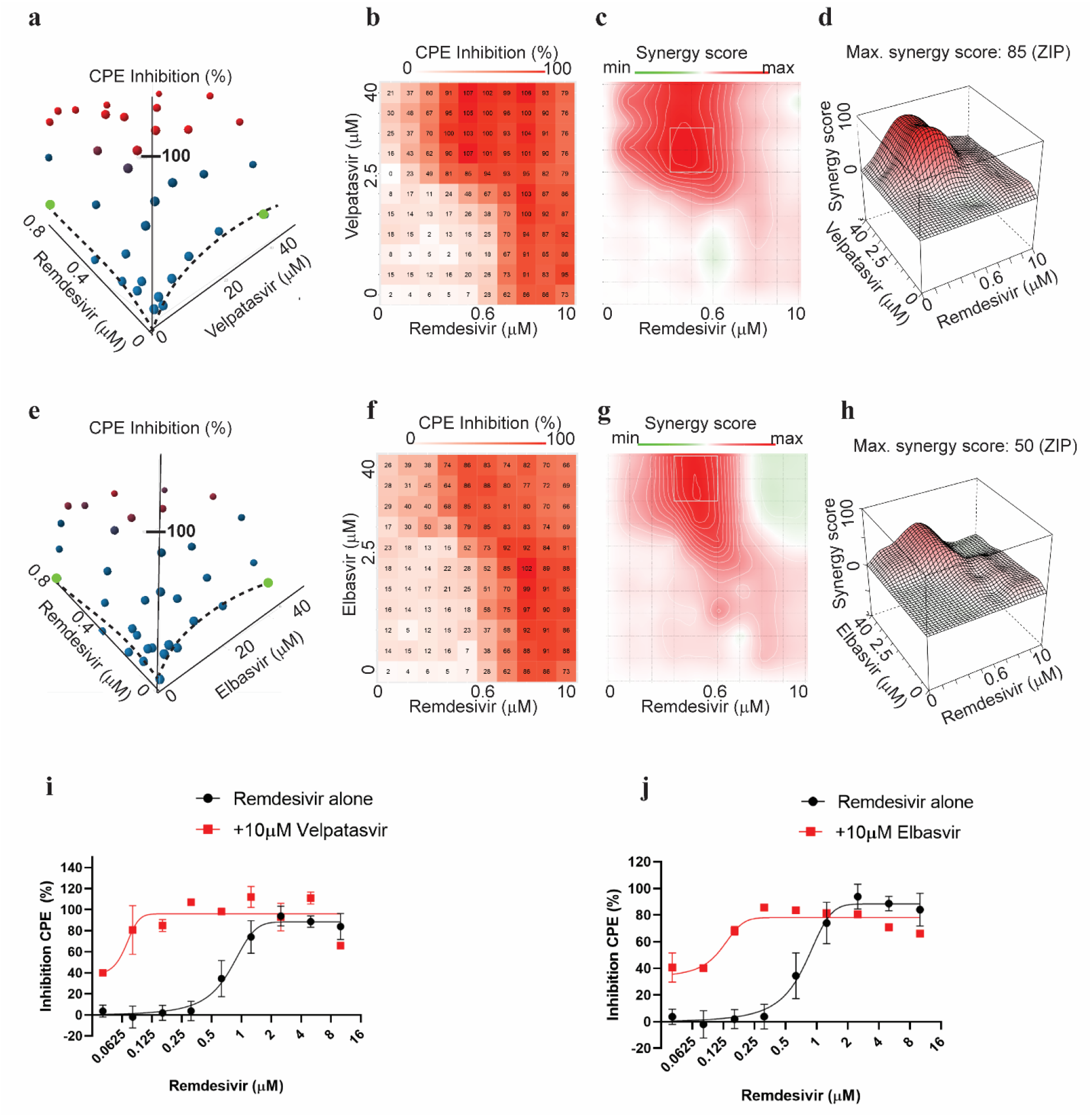
Synergy of direct-acting HCV antivirals velpatasvir (a)-(d) and elbasvir (e)-(h) with Remdesivir in Calu-3 cells infected with SARS-CoV-2. **(a)** Three-dimensional plot showing synergy of combinations of Velpatasvir (x-axis, up to 40μM) and remdesivir (y-axis, up to 0.6μM). Z-axis indicates CPE Inhibition (%). Marker colored using a gradient from blue (0% CPE inhibition) to red (100% CPE inhibition). Green – highest concentration of velpatasvir and remdesivir alone, reaching only ∼20% Inhibition of CPE. Dashed line indicates dose response results of remdesivir and velpatasvir alone, respectively. **(b)** Two-dimensional representation of dose response interaction matrix. X-axis – Remdesivir (up to 10μM), y-axis: Velpatasvir (up to 40μM). Color gradient indicates Inhibition of CPE (%); white – 0%, red-100%. **(c)** Topographic two-dimensional map of synergy scores determined in synergyfinder ^33^ from the data in (a) and (b), axes as in (b), color gradient indicates synergy score (red – highest score). **(d)** Three-dimensional surface plot representing synergy score (z-axis) for each compound combination. X-axis: remdesivir up to 10μM, y-axis: velpatasvir up to 40μM. **(e), (f), (g), (h)** as in (a), (b), (c), (d) but with elbasvir instead of velpatasvir. **(i)** Dose response of remdesivir alone (black) and in combination with 10μM velpatasvir (red). **(j)** Dose response of remdesivir alone (black) and in combination with 10μM elbasvir (red).

Out of this list of strong candidates for remdesivir combination therapy, we prioritized velpatasvir, elbasvir, dabrafenib, cilostazol and nimodipine for detailed characterization based on the strength of the synergistic effect, mechanism of action, safety profile and the likelihood of clinical usefulness in context with best practices for Covid-19 treatment. Velpatasvir and elbasvir are HCV antivirals targeting nonstructural protein 5 (NS5A), a replication co-factor. Dabrafenib is a B-raf inhibitor used for melanoma chemotherapy, with an acceptable safety profile; B-raf inhibitors have been shown to have antiviral effects but have not been reported in context of SARS-CoV-2 ^29,34^. Cilostazol is a widely prescribed, generically available PDE3-Inhibitor, used to prevent stroke and treat intermittent claudication ^35^. Nimodipine is a generically available Calcium channel blocker used to treat hypertension with a favorable safety profile, acting on one of the druggable pathways enriched in the screen (Figure 2) ^36^.

The strongest candidate was velpatasvir (Figure 3), with a maximum synergy score of 87. On its own, 40μM velpatasvir inhibited SARS-CoV-2 replication by less than 20%, as did remdesivir below 0.6μM (Figure 3a, dashed lines/green markers). In combination, 100% CPE inhibition was reached. This was also observed for combinations of submaximal concentrations (Figure 3b). Statistically significant synergy was apparent for combinations from 1μM of velpatasvir and 0.07μM of remdesivir upwards, with a maximum being reached for combinations above 2.5μM velpatasvir and 0.3μM remdesivir (Figure 3c, 3d). The presence of 10μM velpatasvir shifts the EC50 for remdesivir from ∼1μM to 50nM, a 20-fold difference (Figure 3i, red). We also found another HCV NS5A inhibitor, elbasvir, to show synergy in combination with remdesivir. Elbasvir increased inhibition of CPE exclusively when remdesivir was present (Figure 3f), with a synergy score of 50 (Figure 3g, 3h) and measurable effects at concentrations as low as 5μM elbasvir and 0.2μM remdesivir (Figure 3f, 3g, 3h). In presence of 10μM elbasvir, the EC50 for remdesivir was shifted >10-fold, from 0.7μM to about 65nM (Figure 3j). Dabrafenib, cilostazol and nimodipine showed maximum synergy scores of 50, and close to 100% Inhibition of CPE (Extended Data Figure ED3a, ED3b – dabrafenib, ED3c, ED3d – cilostazol, ED3e, ED3f -Nimodipine).

## Efficacy of clinically used coformulations

As both velpatasvir and elbasvir are only available co-formulated with other antivirals, we tested the combinations used in the marketed drug combinations Epclusa (velpatasvir 100mg/sofosbuvir 400mg, Gilead) and Zepatier (elbasvir 50mg/ grazoprevir 100mg, Merck) in the dose response interaction matrix in SARS-CoV-2-infected Calu-3 cells. For Epclusa, velpatasvir and sofosbuvir were added in 1:4 ratio as in their commercial coformulation. Sofosbuvir alone (up to 40μM) showed very little synergistic effect with remdesivir (Figure 4a), velpatasvir alone (up to 10μM) reproduced the synergy observed previously (Figure 4b), and the combination of velpatasvir/sofosbuvir (up to 10/40μM, respectively) increased synergy with remdesivir further, enhancing activity of previously inactive remdesivir concentrations. The combination remdesivir/Epclusa shifts the EC50 value of remdesivir ∼25-fold, to 37nM (Figure 4c, 4d). Also for Zepatier, the triple combination (elbasvir, grazoprevir, remdesivir) showed stronger synergy with remdesivir than elbasvir alone, shifting the EC50 value of remdesivir ∼20-fold, to about 50nM at 10uM elbasvir/grazoprevir (Figure 4g, 4h). Thus, commercially available drug combinations targeting HCV NS5A protein showed the strongest synergy with remdesivir in inhibiting SARS-CoV-2 to date.

**Figure 4.**
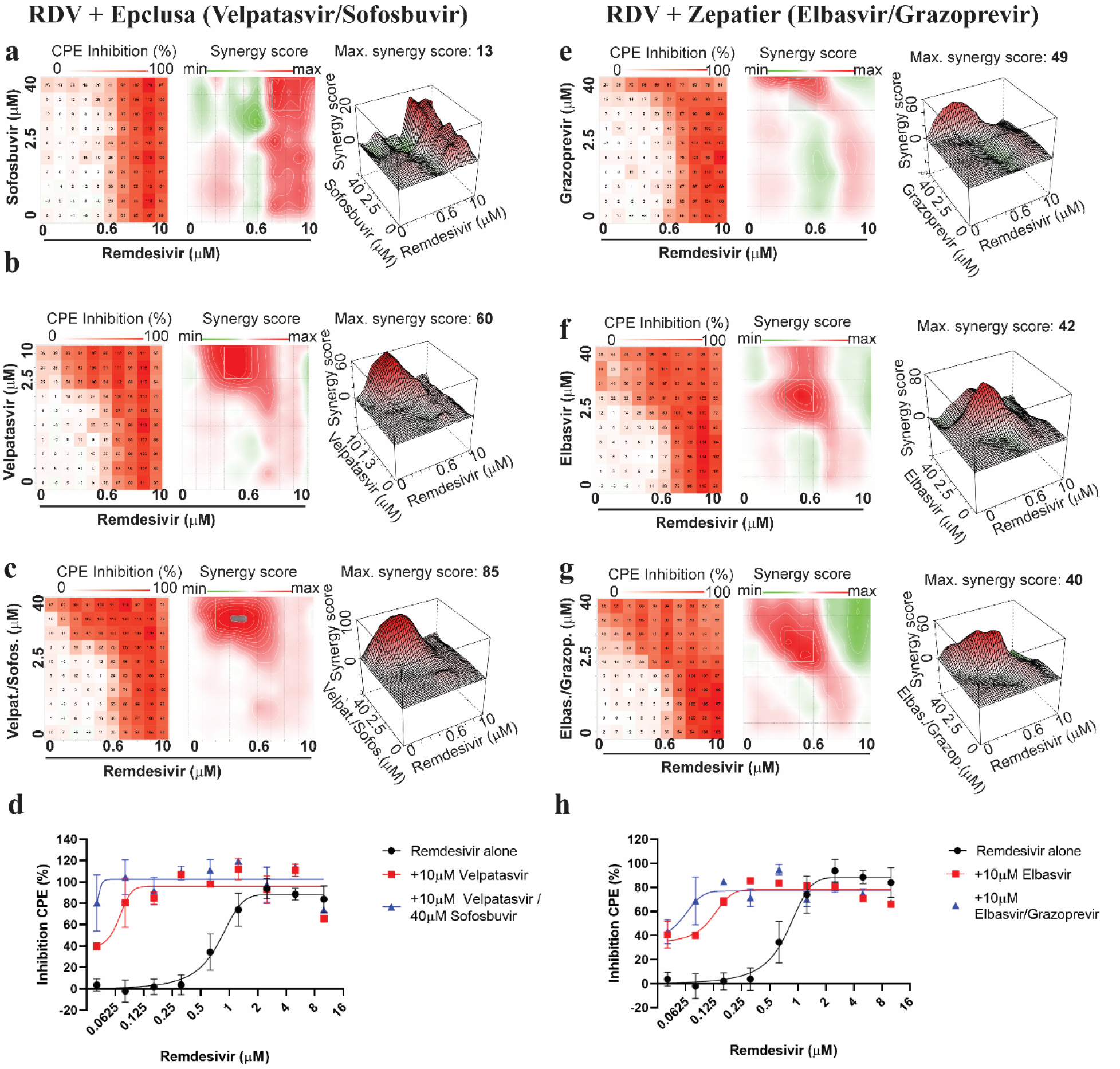
Velpatasvir and elbasvir are more effective in enhancing remdesivir activity when used in their commercially available co-formulation with sofosbuvir and grazoprevir. All experiments shown are in Calu-3 cells infected with SARS-CoV-2.**(a)** Left panel: Two-dimensional representation of dose response interaction matrix. X-axis – Remdesivir (up to 10μM), y-axis: Sofosbuvir (up to 40μM). Color gradient indicates Inhibition of CPE (%); white– 0%, red-100%. Middle panel: Topographic two-dimensional map of synergy scores determined in synergyfinder ^33^ from the data in (a), axes as in (b), color gradient indicates synergy score (red – highest score). NB coloring scheme and z-axis autoscales to the highest value observed, inflating small changes for weak compounds such as sofosbuvir. **(b)** As in (a), but remdesivir combined with velpatasvir (up to 10μM); **(c)** as in (a) but remdesivir combined with both velpatasvir (up to 10μM) and sofosbuvir (up to 40μM); axis indicates sofosbuvir concentration only, Velpatasvir is 4x lower. **(d)** Dose response of remdesivir alone (black) and in combination with 10μM velpatasvir (red) or 10uM velpatasvir / 40uM sofosbuvir (blue). **(e)** as in (a), but remdesivir combined with grazoprevir (up to 40μM). **(f)** as in (a), but remdesivir combined with elbasvir (up to 40 μM). **(g)** as in (a), but remdesivir combined with both elbasvir and grazoprevir (both up to 40μM). **(h)** Dose response of remdesivir alone (black) and in combination with 10μM elbasvir (red) or 10uM elbasvir / 10uM grazoprevir (blue).

## Orthogonal validation of prioritized anti-SARS-CoV-2 drug combinations

We next assessed viral infectivity in a tissue culture infectious dose 50 (TCID50) assay, which determines the titer of infectious viral particles after compound treatment. In this experiment, Calu-3 cells infected with SARS-CoV-2 were treated with remdesivir by itself (EC15) or in combination with velpatasvir, sofosbuvir, elbasvir, grazoprevir, velpatasvir/sofosbuvir, elbasvir/grazoprevir, as well as with dabrafenib, cilostazol and nimodipine. The TCID50 assays confirmed results seen in the screening assay (Figure 5a, 5e) -on its own, remdesivir at its EC15 had only modest effects and velpatasvir or sofosbuvir had no significant effect; yet in combination, remdesivir/velpatasvir/sofosbuvir reduced the titer of infectious viral particles by ∼1500-fold (Figure 5b). Similar results were observed for elbasvir – viral titer was reduced ∼1500-fold in combination of elbasvir and remdesivir, with no or little effect of single agent treatment (Figure 5f). Consistent with earlier results, the co-formulated combination of grazoprevir and elbasvir was synergistic with remdesivir (Figure 5f; Figure 4d). Dabrafenib, cilostazol and nimodipine also reduced viral titer in presence of remdesivir (Figure ED3h).

**Figure 5.**
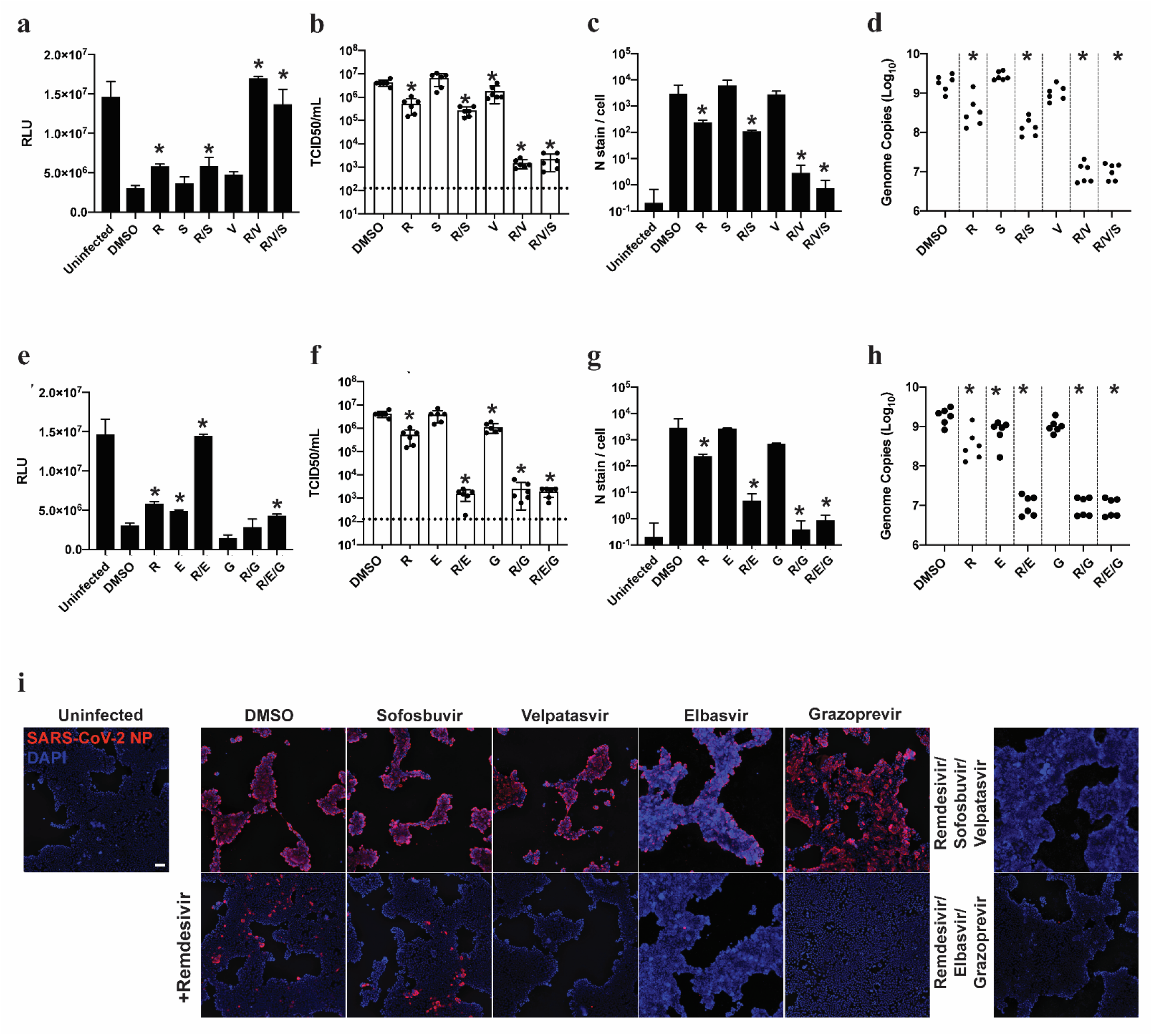
FDA-approved compounds synergize with low dose remdesivir to inhibit SARS-CoV-2 replication in orthogonal assays. **(a) (e)** Cell-titer Glo assay measuring ATP content of viable cells 96h post-infection (hpi) in human lung epithelial Calu-3 cells infected with SARS-CoV-2 (MOI=0.05). Drug was added at 40μM except remdesivir, which was added at 0.625μM (∼EC15), and velpatasvir, which was added at 10μM to maintain the ratio of 1:4 in dosing with its combination sofosbuvir. n=3, error bars indicate Standard deviation. Asterisk indicates statistical significance with p<0.05 relative to DMSO control. R: remdesivir; V: velpatasvir; S: sofosbuvir; E: elbasvir; G: grazoprevir **(b) (f)** infectious virus particle titer leading to 50% of cell death in Vero-E6 cells (TCID50) was determined from the supernatants of (a) and (e) 24hpi; other conditions as in (a) and (e). Dotted line indicates limit of detection in the assay. **(c) (g)** infection was quantified by direct visualization of virus particles by immunofluorescence assay, detecting number of SARS-CoV-2 nucleoprotein (N stain) 48hpi per infected cell. Other conditions were as in (a) and (e). For remdesivir/velpatasvir/sofosbuvir, results were not statistically significantly different from uninfected control cells (p=0.18), as was the case with remdesivir/elbasvir/grazoprevir (p=0.07) **(d) (h)** RT-qPCR quantifying SARS-CoV-2 genome equivalents of Calu-3 cells treated with the indicated drug combinations and infected with SARS-CoV-2 at MOI of 0.05 for 48hpi. **(i)** Representative images from (c) and (g), Calu-3 cells infected with SARS-CoV-2 48hpi. Scale bar corresponds to 100μm.

Similar results were obtained when we quantified infected cells by immunofluorescence microscopy. We treated Calu-3 cells with remdesivir and compound, infected cells with SARS-CoV-2, and stained for nuclei (DAPI) and SARS-CoV-2 Nucleocapsid Protein (N-protein, NP; Figure 5c, 5g, 5i). While EC15 concentrations of remdesivir had little effect on viral replication, as indicated by distinct N-protein staining, its combination with velpatasvir, elbasvir, velpatasvir/sofosbuvir, elbasvir/grazoprevir, dabrafenib, cilostazol or nimodipine strongly reduced the number of infected cells (Figure 5c, 5g, 5i and ED3h). In fact, cells treated with the commercial HCV antiviral combinations were statistically not significantly different from uninfected cells (Figure 5c, 5g, 5i; RDV/velpatasvir/sofosbuvir p=0.18, RDV/elbasvir/grazoprevir p=0.07). Analyzing viral genome copy number from the supernatant of infected cells by qRT-PCR further confirmed the drastic effects of combining HCV antivirals with remdesivir in blocking SARS-CoV-2 replication (Figure 5d, 5h and ED3j). We conclude that the approved HCV antiviral medications Epclusa (velpatasvir/sofosbuvir) or Zepatier (elbasvir/grazoprevir) are strongly synergistic with remdesivir in blocking SARS-CoV-2 replication, significantly reducing viral load of infected cells.

## Discussion

Combination therapy cures Hepatitis C and enables long-term HIV suppression without significant development of resistance. Such therapy is highly desirable for the SARS-CoV-2 pandemic, but typically takes more than 10 years to develop. Here, we identify 20 FDA-approved compounds that have potential to treat SARS-CoV-2 by improving efficacy of remdesivir. Most strikingly, several HCV NS5A-inhibitors showed strong synergy with remdesivir in SARS-CoV-2 infected cells, resulting in drastically reduced viral load in treated cells. The intracellular concentration of the active remdesivir metabolite in the human lung is estimated to be between 4-10μM, close to its 7μM IC50 and way below the 18μM IC90 that would be needed to fully inhibit the virus ^19^. Due to systemic toxicity, remdesivir cannot be dosed higher ^19^; the recent inhalation trials aim to increase lung concentration by changing route of administration. The 25-fold shift in potency reported in this study could move the IC90 from 18μM to ∼0.7 μM in the example above, well below the estimated intracellular concentration of 4-10μM, putting virus eradication within reach. In addition, our proposed combinations could extend the reach of the available remdesivir supply, almost in its entirety stockpiled by the US government. This could allow treatment of more than 5 million Covid-19 patients with the doses manufactured in September alone (∼230000 treatment courses). Therefore, we suggest that the combinations identified in this study should be fast-tracked for in vivo studies and clinical evaluation, for example in inhalation trials.

The molecular target of velpatasvir and elbasvir, NS5A, is a component of the HCV membrane-bound replication complex, with additional roles in virion assembly and modulation of host cell physiology ^37-40^. This complex is also targeted by remdesivir, which might thus represent an example of partial inhibition at two independent binding sites of the same complex synergizing to achieve full inhibition. Due to the similarity between the HCV and coronavirus replication machineries, HCV replication inhibitor Epclusa (velpatasvir/sofosbuvir) was prioritized as a repurposing candidate early in the pandemic ^41^, but did not inhibit virus growth when administered as single agent ^28^. It is possible that the interaction with the SARS-CoV-2 machinery is on its own insufficient to interfere with virus replication, but together with remdesivir, which inhibits the RNA-dependent RNA-polymerase, cripples the virus.

The synergistic effects could be further enhanced by pharmacokinetic interactions. NS5A inhibitors can inhibit the membrane transporters P-gp, BCRP, OATP1B1 and OATP1B3 ^42^. Those transporters reduce intracellular drug concentrations, and their inhibition could increase the apparent potency of remdesivir. However, analysis of known transporter inhibitors in our compound collection reveals only modest enrichment of OATP1B1 inhibitors, and no enrichment for P-gp, BCRP, and OATP1B3 inhibitors (Figure ED4). This indicates that the observed synergy is likely due to the on-target effects of velpatasvir and elbasvir on the viral replication machinery.

In addition to HCV antivirals, we found 18 more synergistic combinations between remdesivir and approved drugs with a favorable safety profile and a wide range of pharmacokinetic properties, including dabrafenib, nimodipine and cilostazol. We also identified the well tolerated and widely used steroids budesonide and meprednisone as showing robust synergy with remdesivir, supporting the notion that steroids can have direct antiviral effects (Figure ED2) ^43,44^. These findings open up the possibility to find dual-action steroid-remdesivir combinations that have antiviral effects early in the infection and exert immunomodulatory effects, as achieved with dexamethasone, later. Synergy with remdesivir was also observed for compounds modulating calcium channel and proton pump activity, consistent with well-established modulation and exploitation of host cell calcium signaling during infection ^30,31,45^. We identified as synergistic with remdesivir the generic calcium-channel blockers nimodipine and nifedipine which are widely used as anti-hypertensives and have an excellent safety profile ^36^. Omeprazole sulfide is a metabolite of omeprazole (Prilosec), an over-the-counter proton pump inhibitor to treat reflux, that has been identified before as enhancing the effect of remdesivir on SARS-CoV-2 ^46^. Interestingly, several proton pump inhibitors have been strong hits in other SARS-CoV-2 repurposing screening campaigns^47^.

Taken together, our study leveraged combinatorial screening to discover compounds synergistic with antiviral remdesivir. We identify 20 promising combinations between remdesivir and approved drugs with a favorable safety profile and a wide range of pharmacokinetic properties. Among these, combining remdesivir with the HCV NS5A inhibitor combinations Epclusa (velpatasvir/sofosbuvir) and Zepatier (elbasvir/grazoprevir) increased remdesivir potency 25-fold and practically eliminated SARS-CoV-2 from infected cells, identifying these combinations as strong candidates for fast-tracked clinical evaluation in Covid-19 patients.

## Figure legends

**Extended Data Figure 1.**
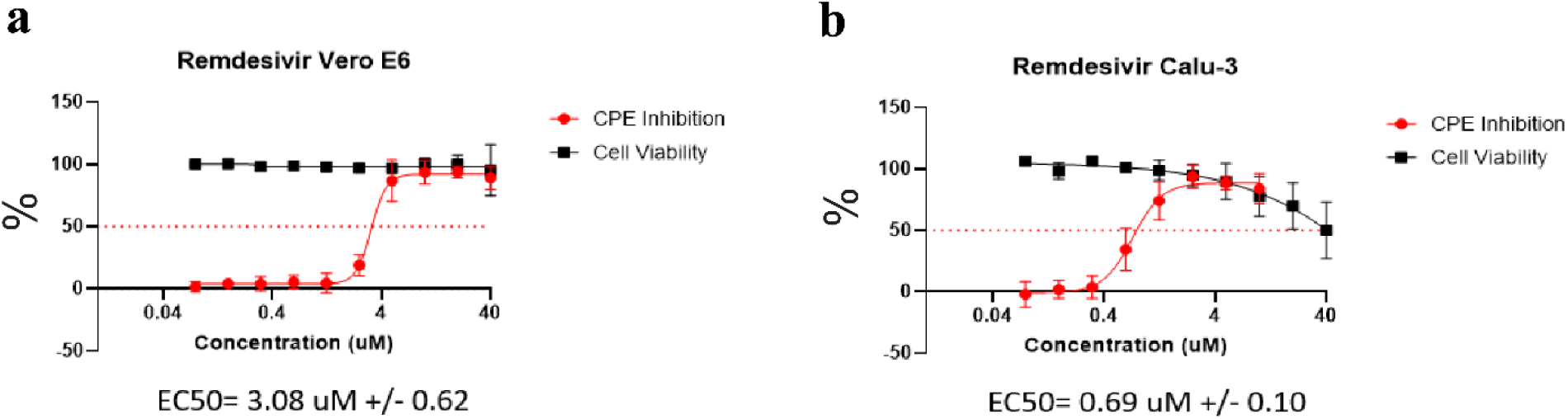
Remdesivir dose response curves averaged across all experiments in 384well. assays in Vero E6 (a) and Calu-3 cells (b). Error bars indicate standard deviation. CPE inhibition (%) in SARS-CoV-2 infected cells in red, Cell viability (%) in uninfected cells in black.

**Extended Data Figure 2.**
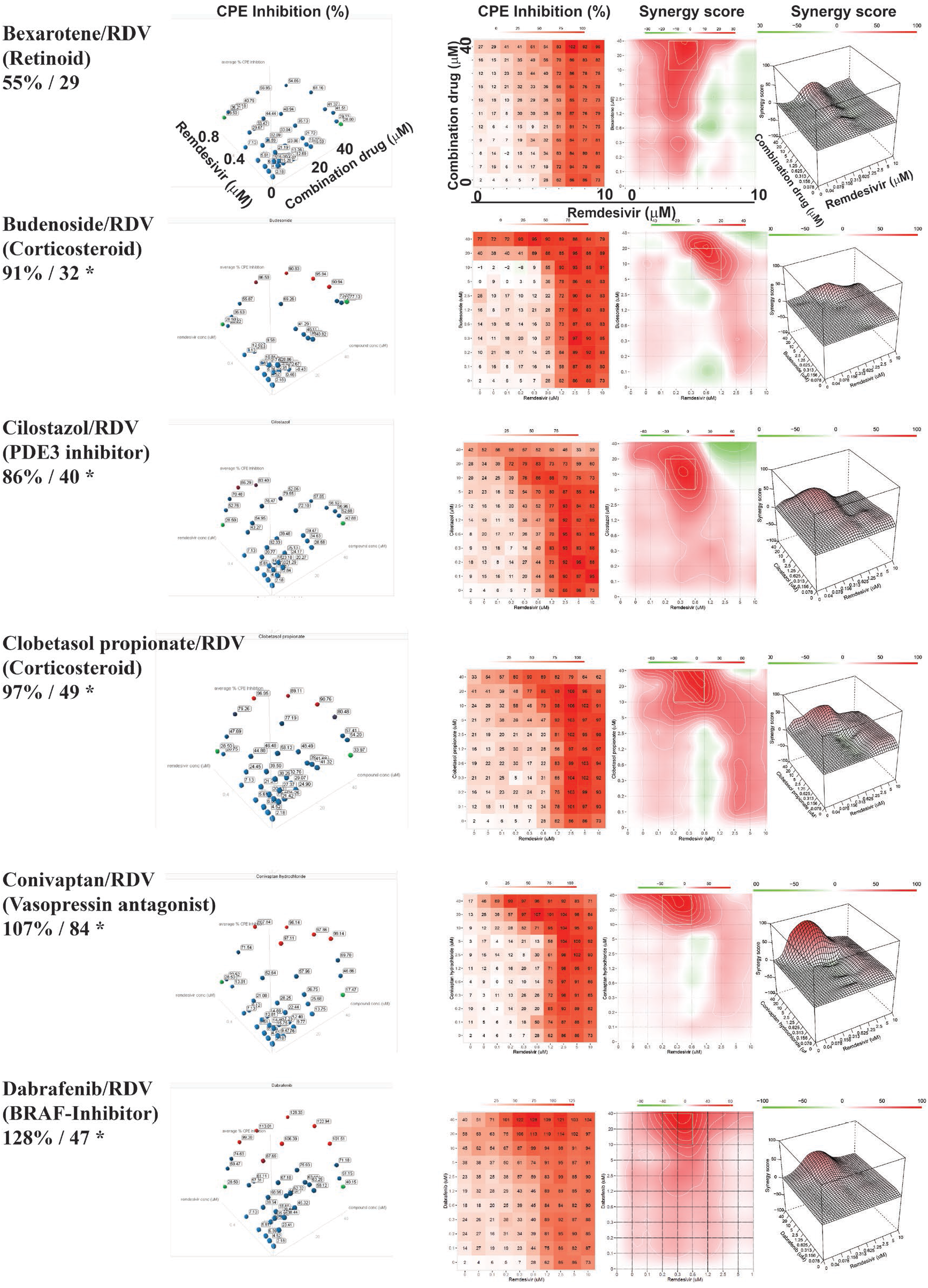
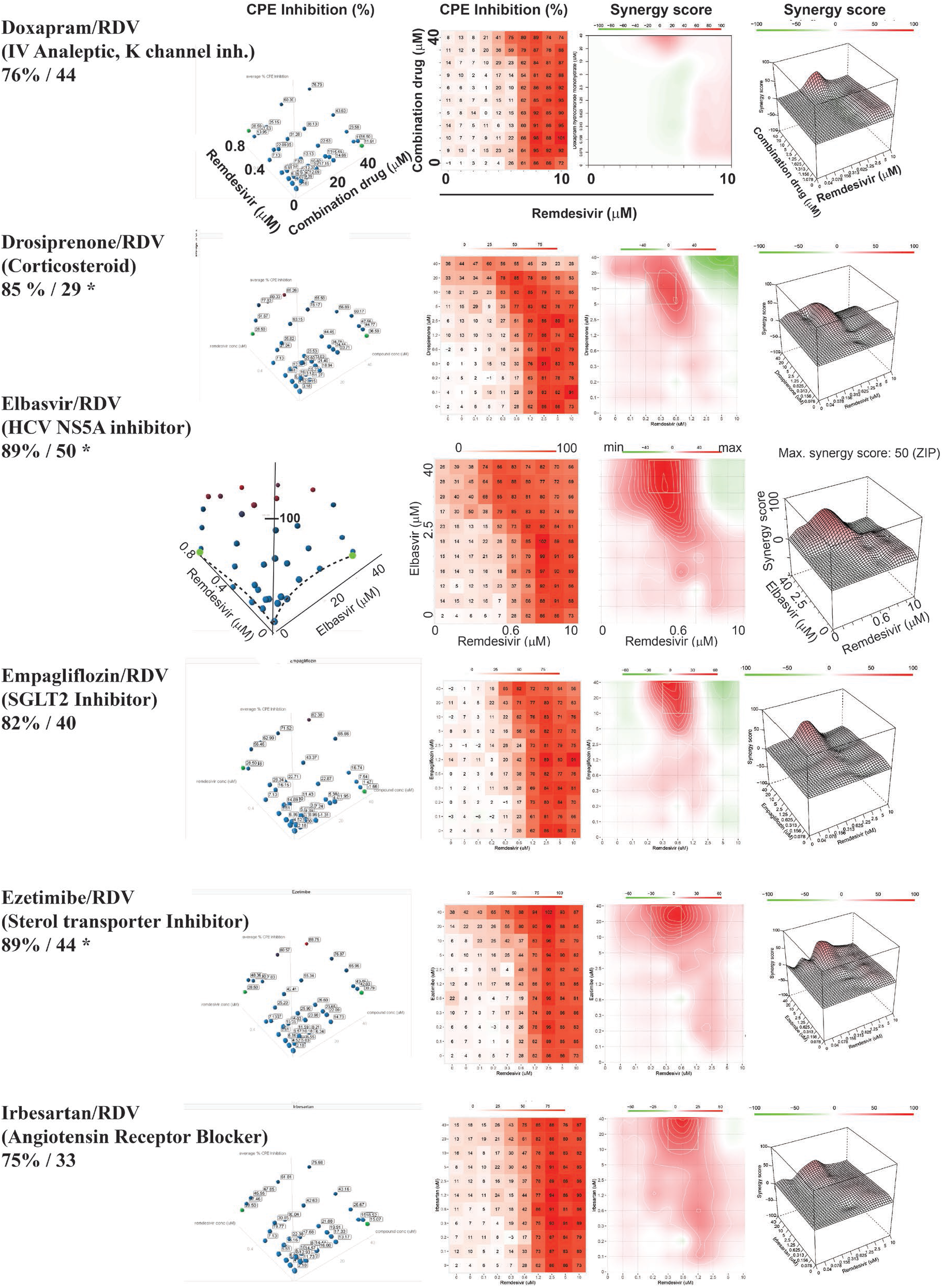
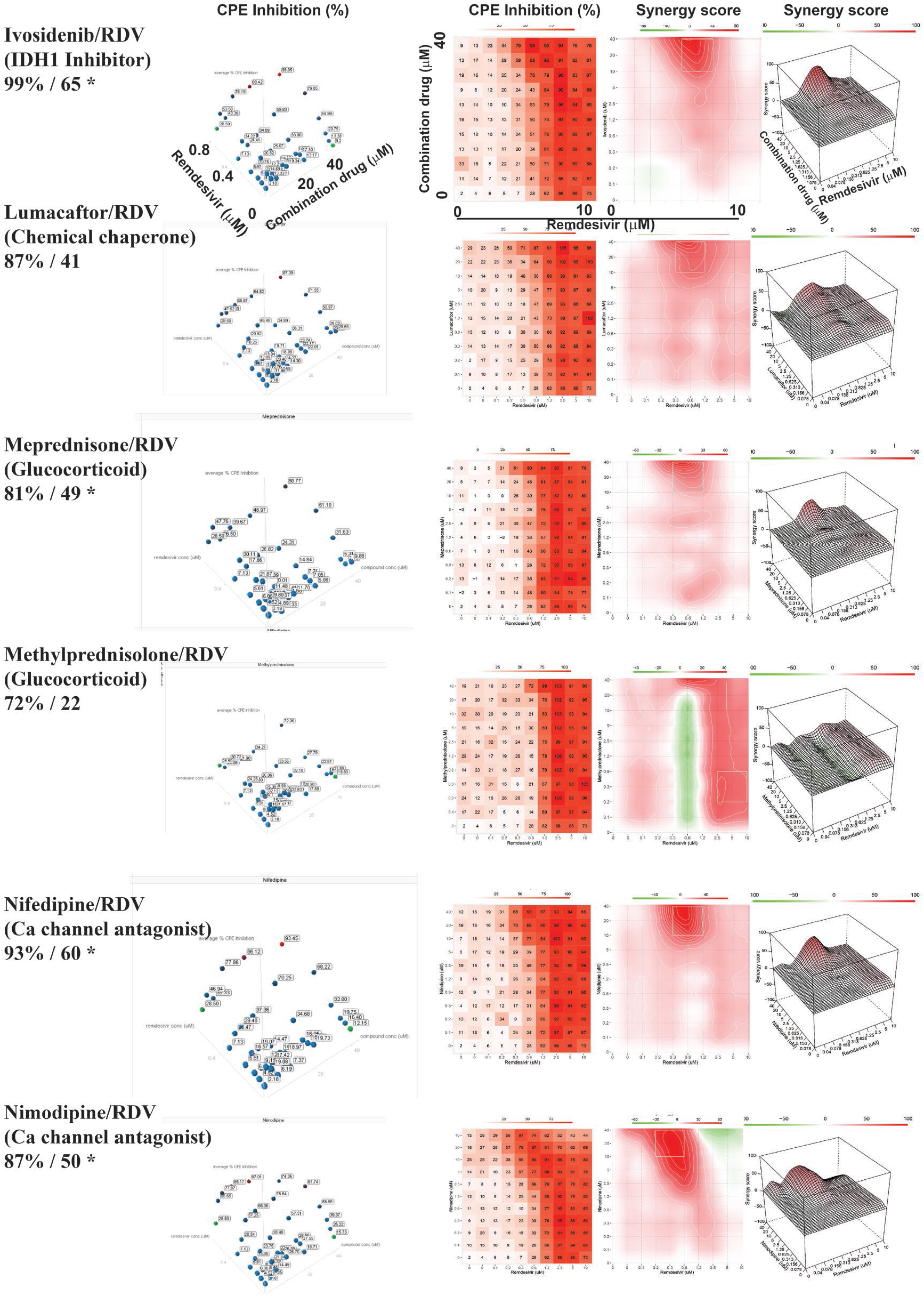
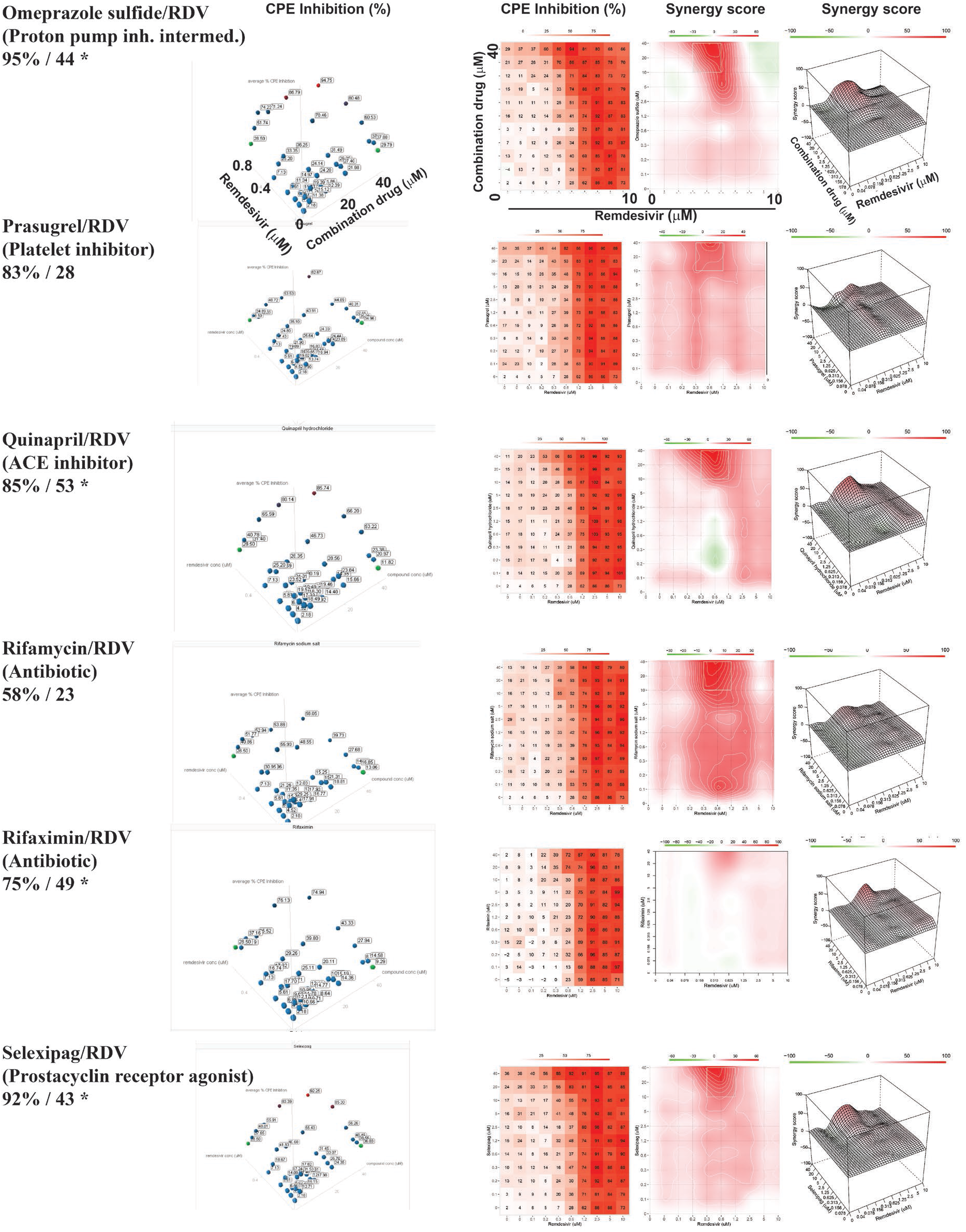
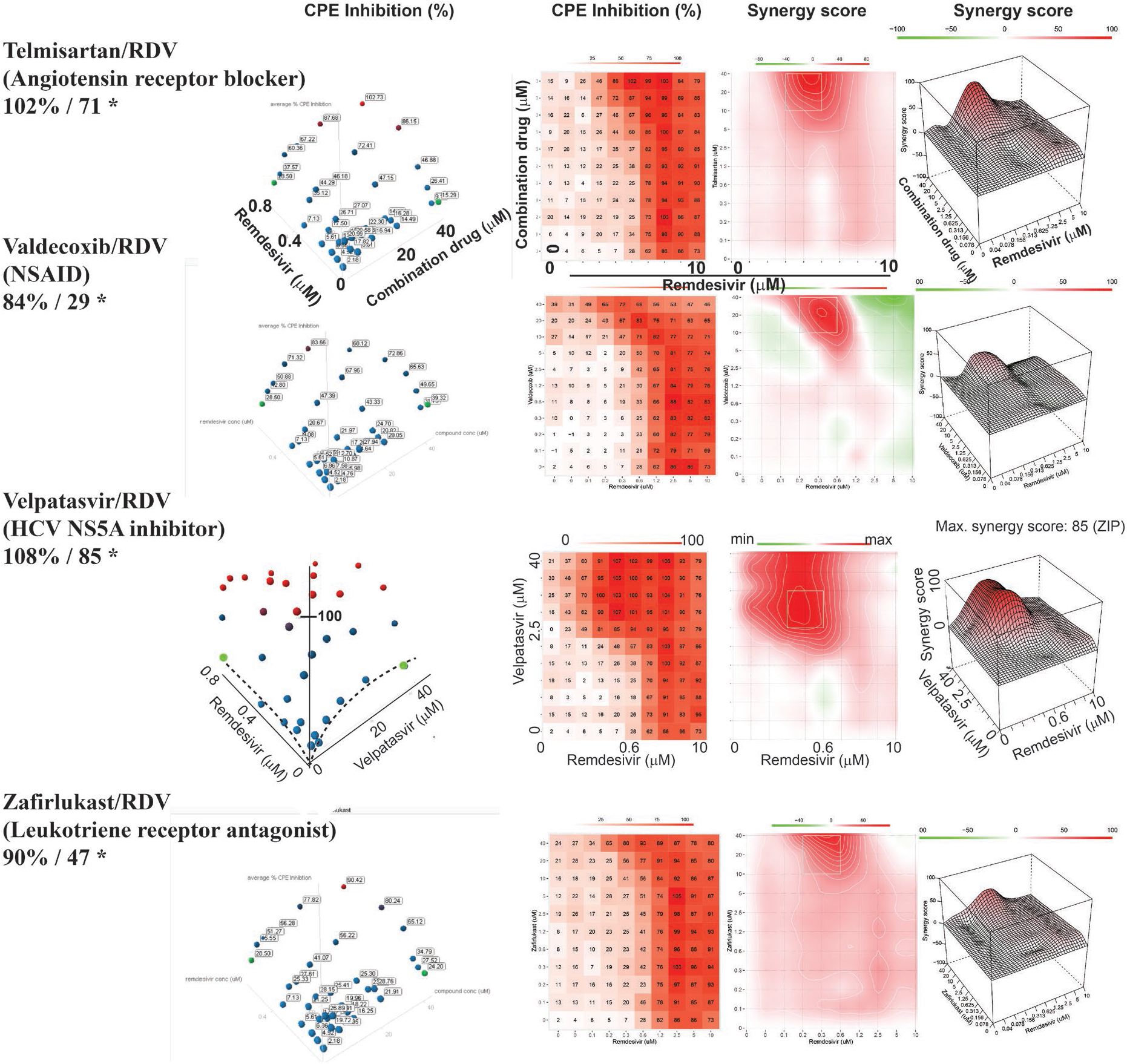
Dose response interaction matrix / synergy analysis for all dose response compound combinations. **Left panel:** Three-dimensional plot showing synergy of combinations of labeled compound (x-axis, up to 40μM) and remdesivir (y-axis, up to 0.6μM). Z-axis indicates Inhibition of CPE (%). Markers are labeled with their respective Inhibition of CPE value; coloring gradient from blue (0% CPE inhibition) to red (100% CPE inhibition). Green – highest concentration of labeled compound and remdesivir alone. Maximum %CPE inhibition and maximum synergy score indicated under drug name. Asterisk indicates 20 prioritized compounds. **Middle left panel:** Two-dimensional representation of dose response interaction matrix. X-axis – Remdesivir (up to 10μM), y-axis: labeled compound (up to 40μM). Color gradient indicates Inhibition of CPE (%); white – 0%, red-100%. **Middle right panel:** Topographic two-dimensional map of synergy scores determined in synergyfinder ^33^ from the data on the left, color gradient indicates synergy score (red – highest score). **Right panel:** Three-dimensional surface plot representing synergy score (z-axis) for each compound combination. X-axis: remdesivir up to 10μM, y-axis: labeled compound up to 40μM.

**Extended Data Figure 3.**
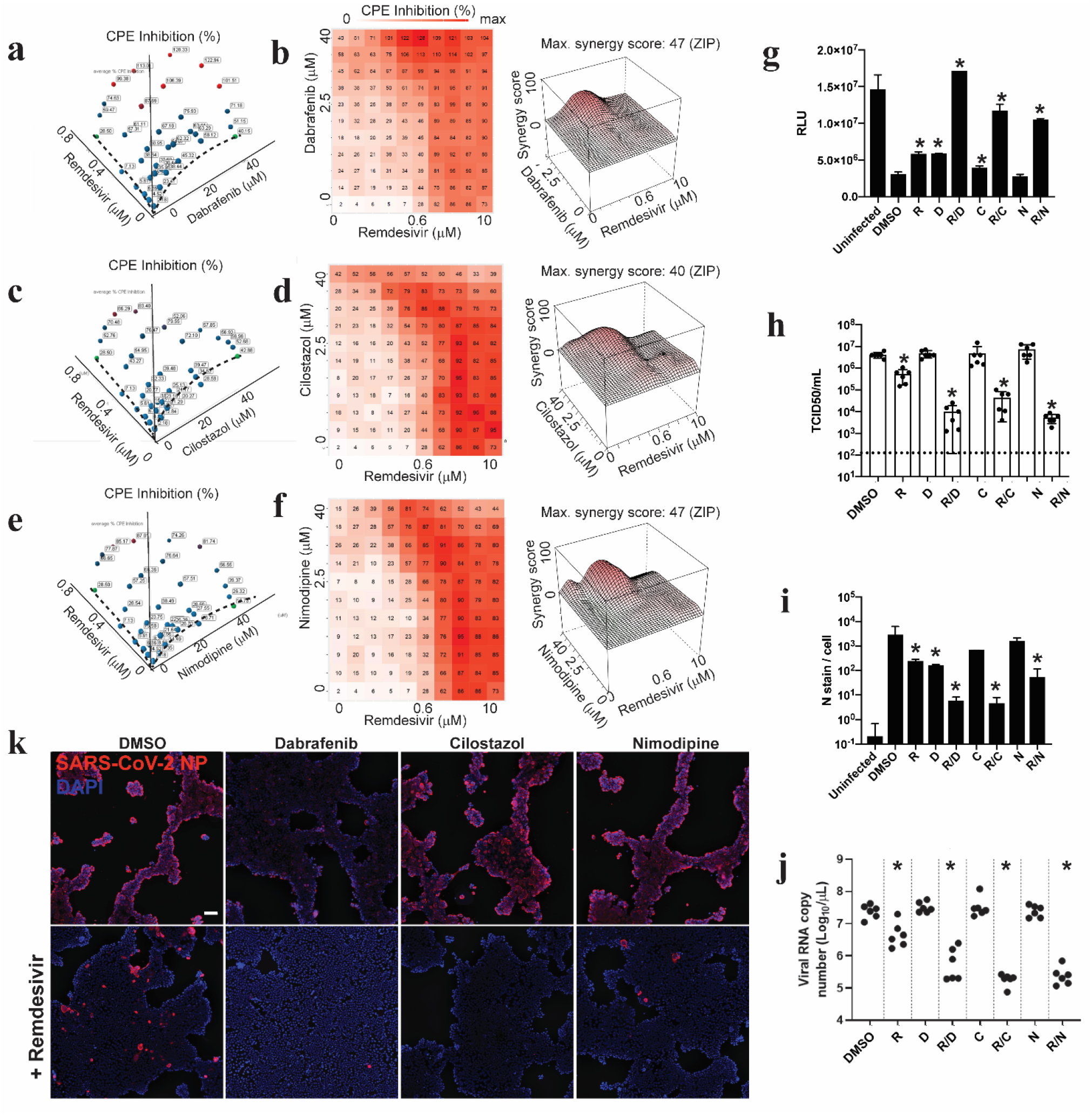
Synergy of dabrafenib, cilostazol and nimodipine with remdesivir in Calu-3 cells infected with SARS-CoV-2. (**a) (c) (e)** Three-dimensional plot showing synergy of combinations of Dabrafenib (a), Cilostazol (c), Nimodipine (e) (x-axis, up to 40μM) and remdesivir (y-axis, up to 0.6μM). Z-axis indicates CPE Inhibition (%). Markers are labeled with their respective Inhibition of CPE value; coloring gradient from blue (0% CPE inhibition) to red (100% CPE inhibition). Green – highest concentration of tested drug and remdesivir alone, reaching ∼20% Inhibition of CPE. Dashed line indicates dose response of each single agent. (**b) (d) (f)** Two-dimensional representation of dose response interaction matrix (left panel) and three-dimensional topographic map of synergy score (z-axis) over dose response matrix. X-axis – remdesivir (up to 10μM), y-axis: Dabrafenib (b), cilostazol (d), nimodipine (f) (up to 40μM). Color gradient indicates Inhibition of CPE (%); white – 0%, red-100%. **(g) (h) (i) (j)** Confirmation of observed activities in orthogonal assays. R: remdesivir; D: dabrafenib; C: cilostazol; N: nimodipine (g) Cell-Titer Glo positive control, (h) TCID50 assay, (i) Immunofluorescence microscopy assay, quantified (j) RT-qPCR assay. Experimental conditions as in Figure 5. **(k)** Representative images from (i), Calu-3 cells infected with SARS-CoV-2 48hpi. Scale bar corresponds to 100μm.

**Extended Data Figure 4.**
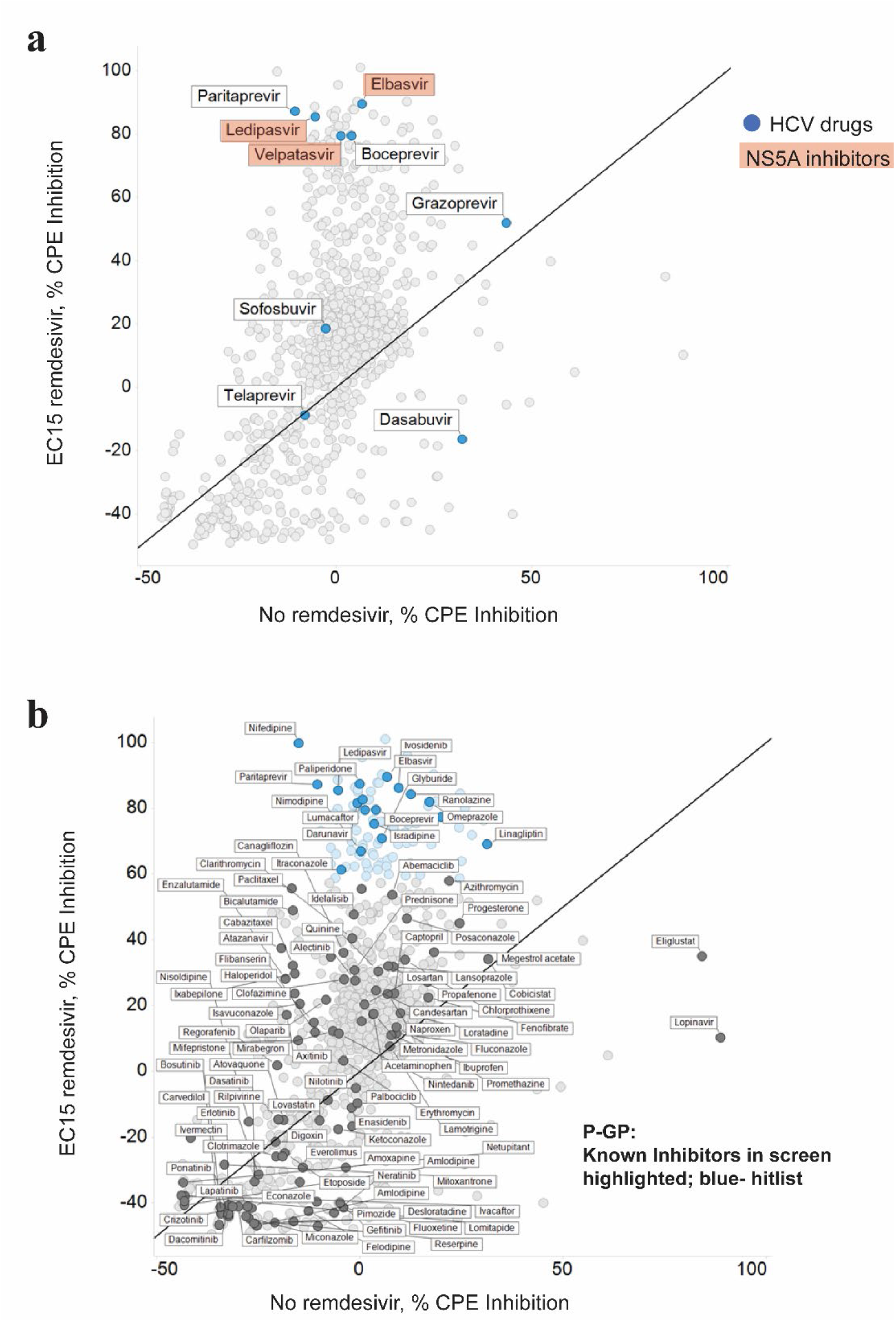
Distribution of NS5A and P-gp inhibitors in the dataset. **(a)** Screening data of all approved drugs in Vero-E6 cells infected with SARS-CoV-2, HCV drugs highlighted in blue, subset of NS5A inhibitors highlighted in red. Axes show inhibition of CPE (%), in absence (x) and presence (y) of EC15 of remdesivir. **(b)** as in (a), but with all known P-gp inhibitors highlighted, showing similar distribution in hitlist (blue) and inactive compounds (gray).

## Methods

### Cells and virus

Vero E6 and Calu-3 cells (Calu-3:ATCC HTB-55; Vero E6: ATCC, CRL-1586) were maintained in high glucose DMEM (Gibco, Waltham, MA, USA) supplemented with 10% FBS (R&D Systems, Minneapolis, MN, USA), 1X GlutaMAX (Gibco, Waltham, MA, USA), and 1X PenStrep (Gibco, Waltham, MA, USA) at 37°C and 5% CO2.

To generate a master viral stock, Vero E6 were plated in T175 flasks (Nunc, Roskilde, Denmark) and allowed to grow to ∼80% confluency before infection with the USA-WA1/2020 strain of SARS-CoV-2 (BEI Resources, Manassas, VA; NR-52281). At 72hpi, dramatic CPE was observed and the flasks were freeze-lysed at -80C. After thaw, lysate was collected and centrifuged at 3000x rpm for 20 minutes to pellet cell debris (Beckman Coulter Allegra X-14R). This procedure was repeated for a second passage working stock with collection at 48hpi and titered by TCID50 assay.

### Compound preparation and drug screening

The FDA-approved drug library containing 1,200 small molecule compounds was stored at 10mM in dimethyl sulfoxide (DMSO) in 384-well master plates (Targetmol, Wellesley Hills, MA). Remdesivir was stored at 10mM in DMSO (T7766, Targetmol). 2500 Vero E6 (12 μl/well) or 10000 Calu-3 (12μl/well) were seeded in 384-well white optical-bottom tissue culture plates (Nunc) with the Multidrop Combi liquid handling instrument (Thermo Fisher Scientific, Waltham, MA). Cells were allowed to adhere and expand, 24 hours for Vero E6 and 48 hours for Calu-3, at 37°C and 5% CO_2_. For the primary screen, confirmation and synergy dose response interaction matrix analysis, compounds were prediluted to 8x final concentration in high glucose DMEM. 3μl compound was transferred from dilution plates using a Cybio Well vario liquid handler (Analytik Jena, Jena, Germany) to cells, leading to a final concentration of DMSO at 0.44% in the assay plate (v/v). Primary screen and confirmation was performed at 40μM compound, dose responses were generated by 2x dilutions starting at 40μM or 10μM. For synergy experiments, EC15(+/-5) of remdesivir was empirically determined and used for each experiment in combination with other drugs as indicated above. Final DMSO was maintained at 0.44 % -0.8% (v/v). Cells were incubated at 37°C and 5% CO_2_ for 1 hour before infection. Viral inoculum was prepared such that the final MOI=0.05 upon addition of 6 μl/well viral inoculum. After complete CPE was observed in DMSO-treated, infected wells 72hpi for Vero-E6 and 96hpi for Calu 3, opaque stickers (Nunc) were applied to plate optical bottoms, and plates were developed with the CellTiter-Glo 2.0 reagent (Promega, Madison, WI) according to the manufacturer’s instructions. For Vero E6 reagent was diluted 1:1 (v/v) in PBS (Gibco, Waltham, MA, USA). Luminescence of developed plates was read on a Spectramax L (Molecular Devices, San Jose, CA). Each plate contained 24 wells uninfected/DMSO treated cells (100% CPE inhibition), and 24 wells infected/DMSO treated cells (0% CPE inhibition). Average values from those wells were used to normalize data and determine % CPE inhibition for each compound well. For duplicate plates, average values and standard deviations were determined. Z’ was determined as described ^48^. Stastical significance was assessed using a two-tailed, heteroscedastic student’s t-test. Measurements were taken from distinct samples unless indicated otherwise. The data was plotted and analyzed with spotfire (Tibco) and GraphPad Prism. Synergy analysis was performed using synergyfinder, using a zero-interaction potency (ZIP) model ^33^.

### GSEA Analysis

Compounds were annotated with targets, pathways and mechanisms of actions using the Center for Emerging and Neglected Diseases’ database and for pharmacokinetic data and transporter inhibition data, the DrugBank database ^42^. Each annotation property was tested for enrichment among the screening hits using the gene set enrichment analysis (GSEA) software as described ^28,49,50^. The compounds annotated for each property were treated as part of the “gene set”. For each set of annotations, the background compound set was defined as the set of compounds annotated for any property. GSEA Preranked analysis was performed using the compounds’ % CPE inhibition from each screen. Compound sets included in the analysis were between 5 and 500 compounds. Enrichment results with p<0.01 and false discovery rate (FDR) q-value <0.1 were considered statistically significant. P-values were generated using a one-sided hypergeometric test^51^.

### Immunofluorescence microscopy analysis (IFA) and RT-qPCR

50000 Calu-3 cells (50μl /well) were seeded in 96-well black optical-bottom tissue culture plates (Nunc). 24 hours post-seeding, drug combinations were added to the cells in 25μl DMEM and incubated at 37°C and 5% CO_2_ for 1 hour before infection. 25μl viral inoculum was added for MOI=0.05. At 48hpi, 75μl supernatant was collected for RT-qPCR analysis. Cells were then washed with PBS, fixed in 4% paraformaldehyde (PFA) in PBS for 15 minutes, washed again with PBS, permeabilized with 0.2% saponin in blocking buffer (2% BSA, 2% FBS in PBS) for 30 minutes at room temperature (RT), and incubated with 1:1000 mouse antibody specific for SARS-CoV-2 nucleocapsid protein (Sino Biological, Beijing, China 40143-MM05) overnight at 4°C in blocking buffer consisting of 2% FBS and 2% BSA. The following day, plates were washed 3X with PBS, incubated with a 1:1000 dilution of goat anti-mouse AlexaFluor647 antibody (Abcam, Cambridge, United Kingdom) and DAPI/Hoechst (Invitrogen) in blocking buffer for 1h at RT, washed again 3x with PBS, fixed in 4% PFA, and replaced in PBS. Plates were fluorescently imaged using an Image Xpress Micro 4 (Molecular Devices). Images were analyzed for N stain per nuclei with the CellProfiler 3.1.9 software (Broad Institute, Cambridge, MA).

For RT-qPCR, 75μl supernatants were collected at 48hpi and inactivated 1:1 in 1X DNA/RNA Shield for RNA extraction and RT-qPCR analysis (Zymo Research, Irvine, CA). RNA was extracted using the QIAamp Viral RNA Mini Kit (Qiagen, Hilden, Germany) according to the manufacturer’s instructions. In brief, 140μl of each sample was mixed with 560μl of Carrier-RNA-containing AVL and incubated for 10min at RT. After addition of 560μl of 100% Ethanol, the samples were spun through columns. The columns were washed sequentially with 500μl of AW1 and 500μl AW2 and RNA was eluted using 50 μl of RNAse free water. RT-qPCR reactions with TaqPath master mix (Thermo Fisher) were assembled following the manufacturer’s instructions.

For a 10 µl reaction, 2.5µl of 4x TaqPath master mix was combined with 0.75 µl of SARS-CoV-2 (2019-nCoV) CDC N1 qPCR Probe mixture (Integrated DNA Technologies, Cat. #10006606, Primer sequences: 2019-nCoV_N1-F: GAC CCC AAA ATC AGC GAA AT; 2019-nCoV_N1-R: TCT GGT TAC TGC CAG TTG AAT CTG; 2019-nCoV_N1-P: FAM-ACC CCG CAT TAC GTT TGG TGG ACC-BHQ1), 3 µl RNA sample, and 3.75 µl water. RT-qPCR was performed on a BioRad CFX96 instrument with the following cycle: 1) 25°C for 1min, 2) 50°C for 15min, 3) 95°C for 2min, 4) 95°C for 3s, 5) 55°C for 30s (read fluorescence), 6) go to step 4 for 44 repetitions. Quantification cycle (Cq) values were determined using the second derivative peak method ^52^. Custom code written in MATLAB (available at https://gitlab.com/tjian-darzacq-lab/second-derivative-cq-analysis) was used to take the numerical second derivative of fluorescence intensity with respect to cycle number, using a sliding window of ± 3 cycles. The peak of the second derivative was fit to a parabola, whose center was taken to be the Cq value ^52^.

### 96 well CellTiter-Glo 2.0 and TCID50 assay

40000 Calu-3 cells (50µl/well) were seeded in 96-well white optical-bottom tissue culture plates (Nunc). 48h post-seeding, drug combinations were added to the cells in 25µl DMEM and incubated at 37°C and 5% CO_2_ for 1h before infection. 25µl viral inoculum was added for MOI=0.05. At 24hpi, 25µl supernatant was saved for TCID50 assay. After complete CPE was observed in DMSO-treated, infected wells 96hpi, opaque stickers (Nunc) were applied to plate optical bottoms, and plates were developed with the CellTiter-Glo 2.0 reagent (Promega, Madison, WI), according to the manufacturer’s instructions. Luminescence of developed plates was read on a Spectramax L (Molecular Devices, San Jose, CA).

To quantify infectious particles secreted by cells in a TCID50 assay, 25μl of supernatant from infected, combination-treated cells was collected at 24hpi/drug treatment and 10-fold serially diluted in DMEM. Each dilution was applied directly to eight wells in 96-well plates (Corning) pre-prepared with Vero E6 cells, then incubated for three days at 37°C and 5% CO_2_. TCID50/mL for each sample was calculated by determining the dilution factor required to produce CPE, including syncytia formation, cell clearing and cell rounding, in half, or 4/8, of the wells. Limit of detection was determined as the concentration of virus resulting in CPE in 50% of the wells treated with the lowest dilution of sample.

## Acknowledgments

The following reagent was deposited by the Centers of Disease Control and Prevention and obtained through BEI resources, NIAID, NIH: SARS-Related Coronavirus 2, Isolate USA-WA1/2020, NR-52281. The authors thank Tomas Cihlar, Hongmei Mo, John Bilello, Gregory Camus at Gilead, and Douglas Fox, Eva Harris, Robert Tjian, Jeffery Cox at UCB and Emmie deWit at NIH for discussions and comments. The authors thank Britt Glaunsinger and Michael Rape at UCB for critical reading of the manuscript. This project was supported by the generosity of Eric and Wendy Schmidt by recommendation of the Schmidt Futures program, through Covid Catalyst funding administered by the Center of Emerging and Neglected Diseases at UC Berkeley, and through Fast Grants (part of Emergent Ventures at George Mason University). EVD is supported by NSF Graduate Research Fellowship DGE-1752814. SBB is an Open Philanthropy Awardee of the Life Science Research Foundation. CDD and TGWG are supported by the Bowes Research Fellows Program N7342, the Siebel Stem Cell Institute W6188, the Jane Coffin Childs Memorial Fund for Medical Research, and the HHMI.

## Author contributions

X.N. generated critical reagents, designed and performed experiments E.W. generated critical reagents, designed and performed experiments, analyzed data E.V.D. generated critical reagents, designed and performed experiments, analyzed data S.B.B. generated critical reagents, designed and performed experiments, analyzed data L.H.Y. performed experiments J.ST. generated critical reagents and computational tools, analyzed data C.D. performed the RNA extraction for all the samples analyzed by RT-qPCR T.G. performed and analyzed the RT-qPCR reactions S.S. oversaw and designed experiments, analyzed data J.S. conceptualization, oversaw and designed experiments, wrote manuscript

## Data availability statement

The authors declare that the data supporting the findings of this study are available within the paper and its supplementary information files.

## Code availability statement

Custom code written in MATLAB for qRT-PCR analysis is available at https://gitlab.com/tjian-darzacq-lab/second-derivative-cq-analysis

